# Preclinical Evaluation of PTK7-Targeted Radionuclide Therapy

**DOI:** 10.1101/2024.11.18.624082

**Authors:** Kim Lindland, Sara Westrøm, Srdan M. Dragovic, Ruth Gong Li, Marion Masitsa Malenge, Betty Ho, Asta Juzeniene, Tina Bjørnlund Bønsdorff

## Abstract

Protein tyrosine kinase 7 (PTK7), a receptor found in tumor-initiating cells, is expressed in various malignancies, including ovarian cancer. While PTK7 has been explored as a target for antibody-drug conjugates, this study is the first to investigate its potential for targeted radionuclide therapy. We developed a murine monoclonal IgG1 antibody (mOI-1) using hybridoma technology and generated a chimeric version (chOI-1) with human IgG1 constant regions. A cell-based screening approach using a library of 6100 cell surface proteins identified PTK7 as the target, confirmed by flow cytometry and surface plasmon resonance analyses. Immunohistochemistry showed strong PTK7 expression in ovarian cancer tissues, and *in vitro* studies demonstrated specific binding and internalization of OI-1 in the ovarian cancer cell line SKOV-3-luc. Biodistribution studies using ^177^Lu-DOTA-mOI-1 injected intravenously in xenograft mice with subcutaneous SKOV-3-luc revealed high tumor uptake and retention. Therapeutic efficacy was assessed by intraperitoneal treatment with ^212^Pb-TCMC-chOI-1 in an intraperitoneal xenograft model, showing significant tumor growth inhibition compared to non-radioactive controls. This study provides the first proof-of-principle for using the PTK7-targeting OI-1 antibody as an antibody-radionuclide conjugate (^212^Pb-labeled), demonstrating its therapeutic potential in a preclinical model of intraperitoneal ovarian cancer. These results support further investigation of OI-1 as a candidate for targeted radionuclide therapy in PTK7-expressing cancers.

## Background

Protein tyrosine kinase 7 (PTK7), a catalytically inactive receptor involved in Wnt signaling, is expressed in various malignancies, including ovarian cancer, and its elevated expression correlates with a poor prognosis (1–5). Interest in PTK7 as a therapeutic target has predominantly focused on its overexpression in tumors to develop strategies, such as antibody-drug conjugates (ADCs) and chimeric antigen receptor T (CAR-T) cells (1, 6–8). Additionally, preclinical evidence suggest that anti-PTK7 antibodies can directly inhibit tumor growth (9) and angiogenesis (10). Despite the interest, there has been no investigation into PTK7 as a target for radionuclide therapy. Our study aims to address this gap by evaluating the potential of PTK7-targeted alpha therapy using ^212^Pb-labeled antibodies, offering a novel approach for treating PTK7-expressing malignancies such as ovarian cancer.

Ovarian cancer, the seventh most common cancer among women worldwide, has poor prognosis and presents significant treatment challenges (11), with a global five-year net survival rate of 30-50% (12). Late-stage diagnosis is a major factor contributing to poor outcomes, as most cases are detected after the cancer has spread within the peritoneal cavity (13). Despite advancements in treatment protocols, including cytoreductive surgery, platinum-based chemotherapy, and targeted therapies, such as bevacizumab and PARP inhibitors, the recurrence rate in patients with ovarian carcinoma exceeds 70% (14–17). This high recurrence rate, particularly due to intraperitoneal (IP) disease spread, underscores the unmet need for novel and more effective therapeutic approaches.

Targeted radionuclide therapy leverages the unique properties of radionuclides to deliver cytotoxic radiation directly to cancer cells, thereby minimizing damage to the surrounding healthy tissues (18). This approach can be broadly categorized as therapies that use alpha and beta emitters, each with distinct characteristics and applications. Targeted alpha therapy (TAT) harnesses the high linear energy transfer (LET) of alpha particles, which deposit a large amount of energy over a short distance (40-100 µm), to effectively eradicate micrometastases while minimizing damage to adjacent healthy tissues, making alpha-emitting radionuclides potent cytotoxic agents well suited for treating micrometastatic diseases (19).

Lead-212, a beta-emitting isotope with a 10.6-hour half-life, is suitable for TAT because of its alpha-emitting progeny, bismuth-212 (60.6-minute half-life), and polonium-212 (0.3-microsecond half-life) (20). The bifunctional chelator p-SCN-Bn-TCMC enables ^212^Pb to bind to specific targeting agents, allowing the antibody-radionuclide conjugate (ARC) to deliver cytotoxic radiation to cancer cells (21). Clinical trials of ^212^Pb-TCMC-trastuzumab for ovarian cancer have shown promising outcomes, with mild and transient adverse events (22, 23).

Our evaluation of the novel mAb OI-1 included target identification, immunohistochemistry, binding and cellular internalization studies, biodistribution analysis using lutetuium-177 (^177^Lu), and efficacy studies using ^212^Pb in a preclinical xenograft ovarian cancer model.

## Material and Methods

### Antibodies

A murine monoclonal IgG1 antibody with an unknown target, designated as mOI-1, was generated using mouse immunization and standard hybridoma technology, as described by Bønsdorff et al. (24). The antibody-producing hybridoma was cultured for the purification of mOI-1 antibody by Diatec Monoclonals (Oslo, Norway). A patent corporation treaty application was filed for the OI-1 antibody and its variants (25). A chimeric version of OI-1, designated as chOI-1, containing murine variable regions, human IgG1, and kappa constant regions, was produced by Genscript (Rijswijk, Netherlands) using recombinant cloning and was expressed in Chinese Hamster Ovary cells. All antibody preparations were >99% pure (SEC-HPLC) with endotoxin levels ≤0.1 EU/mg as determined by the manufacturer. The human IgG isotype control used for *in vivo* experiments (31154, lot WJ3400921) was purchased from Thermo Fisher Scientific (Oslo, Norway). Ultra-LEAF purified human IgG1 isotype control used for *in vitro* experiments was acquired from Nordic Biosite (403502, clone QA16A12, Kristiansand, Norway). The antibodies were labeled using an Alexa Fluor 488 (Alexa 488) Protein Labeling Kit from Thermo Fisher Scientific (A10235, Oslo, Norway) according to the manufacturer’s instructions. Goat anti-human Alexa 488 secondary antibody was procured from Thermo Fisher Scientific (A11013, Lot 2273669, Oslo, Norway).

### Identification of target

Retrogenix Cell Microarray Technology (Charles River [formerly Retrogenix], UK) was used to identify the target antigen of the OI-1 antibody. This involved a multi-phase screening process using a library of transfected HEK293 cells. The microarray technology is based on slides spotted with vectors expressing individual targets as well as ZsGreen for transfection control. The library of target genes encodes the full-length human plasma membrane proteins, cell-surface-tethered human secreted proteins and heterodimeric proteins to be screened, and are reverse-transfected into HEK293 cells. Initially, a pre-screen was performed with slides spotted with expression vectors encoding negative and positive controls (mouse FCGR1, human CD86 and EGFR). The purpose of the pre-screen was to assess transfection efficiency and to evaluate potential non-specific background binding of the mOI-1 antibody to native, untransfected, HEK293 cells. The mOI-1 antibody was tested at concentrations of 2, 5, or 20 µg/mL, alongside antibody controls and added to live transfected cells before fixation. Binding was assessed using an AF647 anti-mIgG H+L detection antibody validated for this system. The following library screen involved over 6100 expression vectors representing human plasma membrane proteins, secreted proteins, and 396 heterodimers arrayed in duplicate. The mOI-1 antibody (20 μg/mL) was screened against two replicates of slide sets in the library screen, and interactions were defined by increased signal intensity compared to the background and analyzed using ImageQuant software (GE Healthcare, Chicago, US). The interactions were categorized as very weak to strong.

Confirmation screen arrays, in which each interaction identified in the library screen along with controls (mouse FCGR1, human CD86 and EGFR), were incubated with 20 µg/mL mOI-1 antibody, murine isotype control, or anti-CD86 CTLA4-mFc. Confirmation screens were performed with antibody incubation on either arrays of live cells before fixation (n=2 slides) or live cells without fixation (n=1 slide). Binding was assessed by fluorescence imaging. A verification of target interactions was performed by flow cytometry using HEK293 cells transfected with expression vectors encoding ZsGreen1 alone or ZsGreen1 in combination with various proteins identified as potential targets in previous steps. Specifically, the transfected proteins included PTK7 (either isoform 1 or isoform 4), ASGR2 (either isoform 1 or isoform 3), GH1, THPO, ROBO2, and CD86, which served as an assay control. The transfections were performed in duplicate to ensure reproducibility. Following transfection, live cells were incubated with either 20 µg/mL of the mOI-1 antibody, 0.2 µg/mL CTLA4-mFc (used as a positive control), or assay buffer alone (negative control).

### Validation of target by SPR analysis to recombinant human PTK7 protein and characterization of affinity

Surface Plasmon Resonance (SPR) analyses of chOI-1 was conducted by Abzena (Cambridge, UK) to validate the target and characterize the affinity. The study utilized two proteins: PTK7 from Acro Biosystems (cat.no: PT7-H52H3, 4313A-21BVF1-14Z) and ROBO2 from Sino-Biological (cat.no: 10310-H08H, Lot LC17JA1720). SPR was performed using multicycle kinetics on a Biacore 8K system using a Protein A capture sensor chip (Cytiva, Cat No. 29127555) with HBS-P+ running buffer containing 1 mg/mL BSA. The analysis temperature was maintained at 25°C with a flow rate of 30 μL/min. The association and dissociation times were set to 240 and 900 seconds, respectively. Chip regeneration was performed using 10 mM glycine-HCl (pH 1.5). chOI-1 was captured at approximately 100 RU, while ROBO2 and PTK7 analytes, after being buffer exchanged into the running buffer, were used in an 8-point, 2-fold dilution series (1.56 nM to 200 nM).

To evaluate and compare the binding characteristics of mOI-1 and chOI-1 antibodies, a second experiment was conducted using multicycle kinetics on a Biacore T200 instrument. The CM5 chip (Cytiva, Cat No. 29104988) was coupled with anti-mouse IgG antibody (Cytiva, cat. BR100838) and anti-human IgG (Cytiva, cat. BR100839) to compare the binding of mOI-1 (murine antibody) to chOI-1 (chimeric antibody) in the same experiment. The experimental conditions as above remained similar except: The regeneration conditions differed between surfaces (3M magnesium chloride for human IgG, 10 mM Glycine-HCl, pH 1.7 for mouse IgG). The analyte, consisting of PTK7, was tested using an 8-point, 2-fold dilution series ranging from 1.56 nM to 200 nM. This experimental design allowed for direct comparison of the binding characteristics of mOI-1 and chOI-1 to PTK7 under identical conditions.

TCMC-conjugated chOI-1 antibodies were tested using a setup similar to that used in the first experiment described above with modifications: Single cycle kinetics (SCK) was used in this experiment. A 5 point 2-fold dilution series of PTK7 from 12.5 to 200 nM without regeneration between each concentration was used, and the association and dissociation times were adjusted to 180 s and 300 s, respectively.

### Immunohistochemical staining

Immunohistochemistry was performed by LabCorp (Cambridge, UK). The test system consisted of cryosections prepared from primary ovarian cancer tissues. mOI-1 and chOI-1 and control antibodies (human IgG1, cat. 02-7102, Thermo Fisher Scientific, UK and mouse IgG1 cat. 02-6100; Thermo Fisher Scientific, UK) were conjugated with Alexa 488 fluorescent dyes to facilitate immunohistochemical detection. Methods were developed to generate a suitable immunohistochemical staining protocol involving titration of test articles and control antibodies at various concentrations (10, 2.5, and 0.625 μg/mL) in ovarian cancer tissue sections. Staining intensity was scored on a scale from 0 to 4, with 0 indicating no staining, 1 indicating weak intensity, 2 indicating mild intensity, 3 indicating moderate intensity, and 4 indicating high-intensity staining that may obscure cellular structures. The staining frequency was graded from -to ++++ based on the percentage of cells showing positive staining, where - indicates no staining, + indicates positive staining in ≤25% of cells, ++ in 26-50% of cells, +++ in 51-75% of cells, and ++++ in ≥76% of cells. The assessment was performed by trained histologists and reviewed by a qualified pathologist when required at LabCorp (UK).

### Cell lines

The human ovarian cancer cell line SKOV-3-luc, obtained from Bioware/Caliper Life Sciences (cat. no. 78425 Hopkinton, MA, USA), and the osteosarcoma cell line OHS (established at the Norwegian Radium Hospital) (26) were used in this study. SKOV-3-luc cells were established from ascites of a patient with ovarian adenocarcinoma (27). OHS cells originated from a patient with osteosarcoma (26). The cells were cultured in cell medium: McCoy’s 5a for SKOV-3-luc (Fisher Scientific, Oslo, Norway) or RPMI 1640 for OHS (Life Technologist Invitrogen, Thermo Scientific, Waltham, MA, USA), supplemented with 10% heat-inactivated fetal bovine serum (FBS) (Fisher Scientific, Oslo, Norway) and 1% penicillin-streptomycin (Fisher Scientific, Oslo, Norway) in an incubator at 37°C with 5% CO_2_. At 80-90% confluence, the cells were propagated and harvested by brief detachment with TrypLE Express (Fisher Scientific, Oslo, Norway) and centrifuged at 1200 rpm for 5 min.

### Measurement of PTK7 expression on SKOV-3-luc ovarian cancer cells

The SKOV-3-luc ovarian cancer cell line was harvested and washed with Dulbecco’s phosphate-buffered saline (DPBS, VWR, Oslo, Norway), and the cell number and viability were determined using a Countess Cell Counter (Invitrogen, Carlsbad, USA). The cells were diluted in flow buffer; DPBS containing 0.5% bovine serum albumin (VWR, Oslo, Norway). For saturation experiments, Alexa 488-labeled chOI-1 antibody was added to 0.2 × 10^6^ cells per tube at concentrations ranging from 0.5 to 100 nM. Cells were incubated at 4°C for 3 h with occasional shaking. Following incubation, the cells were washed twice with 500 µL flow buffer and centrifuged at 1200 rpm for 5 min. All cells were stained with MitoTracker Red CMXRos (Thermo Scientific, Oslo, Norway) for 30 min to specifically label mitochondria in live cells (28). The cells were then washed twice with the flow buffer. The washed cell pellets were resuspended in 100 µL of flow buffer with 1 µM DRAQ5 (Fisher Scientific, Oslo, Norway) added 10 min before sample collection. DRAQ5 is a cell-permeable reagent with high affinity for double-stranded DNA and is used to exclude dead cells (29). The samples were collected in triplicate using a CytoFLEX Flow Cytometer (Beckman Coulter, Indianapolis, USA). The experiments were repeated at least twice on independent days. Data were analyzed using CytExpert version 2.4 (Beckman Coulter, Indianapolis, USA). Gating was performed on singlets, followed by the specific gating of MitoTracker Red CMXRos and DRAQ5 double-positive populations to exclude dead cells. The assay was not corrected for non-specific binding; however, the cells were treated with 0.5% BSA 30 min before the addition of Alexa 488-labaled chOI-1 to reduce non-specific binding. For statistical analysis, the relative median fluorescence intensity (rMFI) was calculated and compared with that of unstained cells using the following formula: (rMFI sample – rMFI unstained) / rMFI unstained. Means and standard deviations were calculated from triplicate samples for each experiment. For the final graphs, averages and standard deviations were calculated from multiple experiments. A one-site binding model;

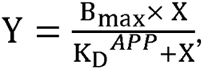

was used to analyze the data, which were plotted as a hyperbolic curve using GraphPad Prism, (GraphPad software, version 10.1.2 La Jolla, USA). Y is the rMFI, B_max_ is the maximum specific binding (max rMFI) and X is the concentration of Alexa 488 labeled chOI-1. This analysis was used to determine the apparent equilibrium dissociation constant (K_D_^APP^) of chOI-1 in the SKOV-3-luc cells.

### Internalization assay

The internalization of chOI-1 in SKOV-3-luc ovarian cancer cells was assessed by flow cytometry using Alexa Fluor 488-labeled antibodies. On the day of the experiment, 0.35 × 10^6^ SKOV-3-luc cells per well were seeded in 24-well plates (Corning, NY, USA) containing 10% complete cell medium. Alexa 488-conjugated chOI-1 antibody (0.25 µg) was added to the cells and incubated at 37°C on an orbital shaker for the indicated time periods (1, 6, and 24 h). Thirty to forty-five minutes before the end of the incubation period, the cells were incubated with MitoTracker Red CMXRos (Thermo Scientific, Oslo, Norway) to specifically label the mitochondria in live cells. The culture medium was removed at designated time points and the cells were washed twice with DPBS. To remove surface-bound antibodies, a stripping buffer containing trypsin (0.01 g/mL in DPBS, pH 8-9, Sigma Aldrich, Oslo, Norway) was added to the cells and incubated for 1 hour, following the method of Lin et al. (30). After stripping, the cells were washed twice with flow buffer. Flow cytometry was performed on both non-stripped and stripped cell samples to assess the change in the Alexa 488 signal under the described conditions for each time point. The cells were analyzed using a CytoFLEX Flow Cytometer (Beckman Coulter, Indianapolis, USA) to determine the degree of receptor internalization compared to the total signal. Surface binding was assessed as relative median fluorescence intensity (rMFI), as previously described. The percentage of internalized signals was calculated as internalized rMFI signal (trypsin-stripped cells) compared to the total-bound rMFI (non-stripped cells), rMFI_Internalized_/rMFI_total_ × 100%.

### Preparation of radionuclides and radioactivity measurements

Lead-212 was produced by emanating ^220^Rn from ^228^Th, sourced from the Oak Ridge National Laboratory (Oak Ridge, USA), or from a ^224^Ra solution, prepared as previously described A simplified single-chamber system was employed, consisting of an inverted 100 mL glass flask with a removable cap containing the radionuclide source. ^228^Th in 1 M HCl was applied to a piece of quartz wool inside the cap that served as the holding material. During decay, ^220^Rn emanated from the quartz wool and was adsorbed onto the interior surface of the flask as ^212^Pb. After 1-2 days of decay, the flask was carefully detached from the source cap and rinsed with 0.1 M HCl to collect the deposited ^212^Pb. The activity of ^212^Pb was measured using a Hidex Automatic Gamma Counter (Hidex Oy, Turku, Finland) with a 60-110 keV counting window, or a Capintec CRC-25R radioisotope dose calibrator (Capintec Inc., Ramsey, NJ, USA) with dial setting 662, a dial setting specifically established for the instrument used (32).

^177^LuCl_3_ solution was obtained from Isotope Technologies Garching (ITG, Germany). ^177^Lu radioactivity was measured using a Cobra II auto-gamma detector (Packard Instrument Company, USA) with a counting window of 15-2000 keV.

### Radiolabeling of antibodies with ^212^Pb and^177^Lu and subsequent IRF measurements

Antibodies mOI-1, chOI-1 and hIgG1 in carbonate buffer were conjugated with 5-20-fold molar excess of S-2-(4-isothiocyanatobenzyl)-1,4,7,10-tetraaza-1,4,7,10-tetra(2-carbamoylmethyl) cyclododecane (p-SCN-Bn-TCMC, TCMC; Macrocyclics Inc, Dallas, USA) in 5 mM HCl (Merck, Darmstadt, Germany) at room temperature for 2 hours. Unbound TCMC was removed by exchanging carbonate buffer with 0.9% NaCl (Merck, Darmstadt, Germany). The extracted ^212^Pb in 0.1 M HCl obtained from the emanation generator was adjusted to pH 5-6 with 5 M sodium acetate (Merck, Darmstadt, Germany) and mixed with TCMC-mAbs with specific activities of 1-50 MBq/mg. The solution was incubated for 30-35 minutes in a Thermomixer (Eppendorf, Oslo, Norway) at 37°C with shaking at 350 rpm and then diluted in a formulation buffer consisting of DPBS with 7.5% (v/v) recombinant albumin (Octapharma, Lachen, Switzerland), 200 mM sodium ascorbate (Sigma-Aldrich, Oslo, Norway), and 1 mM EDTA (Sigma-Aldrich, Oslo, Norway) at pH 7.4, resulting in ^212^Pb-TCMC-mAb. The radiochemical purity of the final products was assessed using chromatography strips from Biodex (Shirley, NY, USA) and gamma counting as previously described. All the ^212^Pb conjugates used in this study had a radiochemical purity of >96%.

For ^177^Lu radiolabeling, mOI-1 antibody was conjugated to p-SCN-Bn-DOTA (Macrocyclics, Dallas, USA) at a of 5 fold molar excess. p-SCN-Bn-DOTA was dissolved in 5 mM HCl and added to the mOI-1 antibody. The reaction mixture was pH-adjusted to approximately 8.5 using 0.1 M carbonate buffer and incubated at 20°C for 2 h with gentle shaking. The unconjugated chelator was separated from the DOTA-conjugated antibody using a centrifuge filtering cartridge (Vivaspin 15R, 50 kDa MWCO, Sartorius Stedim Biotech, Göttingen, Germany), and the buffer was exchanged with 0.9% NaCl. The ^177^Lu solution in 10 mM HCl (ITG, Garching, Germany) was mixed with the DOTA-conjugated mOI-1 antibody in 0.5 M ammonium acetate to adjust the pH to 5.5-6.0 and incubated at 37°C for 15-45 minutes. The radiochemical purity of the products was evaluated using chromatographic strips (Biodex, Shirley, NY, USA). If the purity was below 95%, the conjugate was further purified using a PD10 gel filtration column (GE Healthcare Biosciences, Uppsala, Sweden). The radiochemical purity of the final product was greater than 96%.

The immunoreactive fraction (IRF) assay was performed using cultured SKOV-3-luc cells or cultured or frozen stocks of the high PTK7 antigen-positive OHS. The use of frozen OHS cells simplified the assay procedure while maintaining its effectiveness in quickly and easily evaluating the IRF of radiolabeled antibodies. IRF was measured using one-point binding assays *in vitro* as previously described (33). Briefly, a single-cell suspension, ranging from to 10-15 × 10^6^ cells, was prepared in flow buffer. Triplicate samples, each consisting of 0.2 mL a cell solution, were incubated at room temperature for approximately 60 min with gentle shaking in the presence of 2 ng of radiolabeled mAb. To estimate non-specific binding, two additional samples were preincubated for 15-30 minutes with 20 µg of unlabeled mAb before the addition of the radiolabeled mAb. The total radioactivity of each sample was measured using a Hidex automatic gamma counter or Cobra II auto-gamma detector. Following centrifugation and washing of the cells three times with flow buffer, cell-bound radioactivity was measured in the cell pellets. The fraction of bound mAb was calculated by dividing the amount of cell-bound radioactivity by the total added radioactivity, and immunoreactivity was determined by subtracting the fraction of non-specifically bound mAb from the fraction of bound mAb. The mean IRF and RCP results for each type of radiolabeling are listed in Table S1. The statistical significance of the differences in IRF between the different TCMC:mAb was determined using GraphPad software with a two-sided unpaired t-test, with a significance threshold of p<0.05.

### Determination of number of antigens per SKOV-3-luc cell

A saturation-binding assay was performed to determine the binding characteristics of ^177^Lu-DOTA-mOI-1 to SKOV-3-luc cells. The cells were harvested and resuspended in a flow buffer at a concentration of 10 × 10^6^ cells/ml. Increasing concentrations of ^177^Lu-DOTA-mOI-1 were added to 0.1 ml cell samples in quadruplicate, resulting in final concentrations ranging from 1.6 to 144 nM. Samples were incubated with gentle agitation for 65-95 minutes at 4°C. Non-specific binding was determined by pre-incubating half of the samples with approximately 100 µg/ml unlabeled mAb for 30-60 minutes before adding ^177^Lu-DOTA-mOI-1. After incubation, the cells were separated from the unbound radioligands by centrifugation, and radioactivity in the cell pellet was measured using a Cobra II gamma counter. The values of the equilibrium dissociation constant (K_D_) and the maximum number of binding sites (Bmax) were determined using a one-site binding model, which was plotted as a hyperbolic curve using GraphPad software.

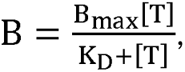

where B is the number of antigen sites per cell and [T] is the total concentration of ^177^Lu-DOTA-mOI-1. The assay was performed in two independent experiments.

### Mass spectrometry analysis of chOI-1 with different TCMC:Ab ratios

An experimental workflow was established and performed by GenScript Biotech (Netherlands) to estimate the number of TCMC conjugated to chOI-1. For proprietary reasons, certain aspects of the mass spectrometry method remain undisclosed by GenScript Biotech. For the experimental workflow, enzyme cleavage was performed using PNGase at 37°C for 4 hours. The PNGase enzyme produced by GenScript was used at a protein-to-enzyme ratio of 50:1. Following enzymatic cleavage, the samples were loaded into an electrospray ionization time-of-flight (ESI-TOF) mass spectrometer (BioAccord System) system from Waters (Milford, Massachusetts, US) for detection. The specific elution procedure used in the ESI-TOF analysis is confidential. For quality analysis, the detected signal was compared to the theoretical molecular weight to determine the number of TCMCs coupled to the protein. Quantity analysis involves quantification of proteins coupled to different TCMCs through signal responses.

### Animals

All animal-related procedures adhered to the Norwegian and EU guidelines for animal research as well as the ARRIVE guidelines (https://arriveguidelines.org/). The study was approved by the Institutional Committee on Research Animal Care of the Department of Comparative Medicine, Oslo University Hospital (Oslo, Norway) and the Norwegian Food Safety Authority (FOTS permit ID: 5734 and 23766). *In vivo* experiments were conducted using female athymic nude mice (Hsd: Athymic Nude-Foxn1^nu^) aged approximately 7-11 weeks, obtained from the Department of Comparative Medicine at the Norwegian Radium Hospital, Oslo University Hospital (Oslo, Norway). In total, 64 animals were included in the study.

### Biodistribution of ^177^Lu-DOTA-mOI-1

Biodistribution experiments were performed using ^177^Lu-DOTA-mOI-1 on 16 athymic female nude mice with subcutaneous SKOV-3-luc xenografts. Mice were injected with 10 × 10^6^ SKOV-3-luc cells in both flanks and tumor growth was monitored 2-3 times weekly using an electronic caliper. Biodistribution experiments commenced when tumor diameters reached 4-15 mm, on day 35 post-inoculation. The maximum tumor size for the mice was defined as 20 mm according to our application to the national animal research authority (FOTS ID 5734). Sixteen mice were intravenously injected with 450 kBq of ^177^Lu-DOTA-mOI-1. Mice were euthanized by cervical dislocation on days 1, 3, 4, and 7 post injection. Selected organs and blood were harvested and their radioactivity was quantified using a Cobra II gamma counter. Statistical analyses were performed using the GraphPad software. Unpaired t-tests were used to determine statistically significant differences in uptake across time points for individual tissues and between tumors and different tissues at each time point. P-values were adjusted using the Holm-Sidak method to account for multiple comparisons, with p_adj_<0.05 set as the significance threshold.

### Therapeutic Efficacy ^212^Pb-TCMC-chOI-1

Tumors were established in 48 female nude mice aged seven weeks by IP inoculation with 5 × 10^6^ SKOV-3-luc cells. Three days after cell inoculation, the mice were randomized into six groups, with eight mice per group: each mouse received an IP bolus of either 1) saline, 2) chOI-1, 3) 211 kBq ^212^Pb-TCMC-hIgG, 4) 384 kBq ^212^Pb-TCMC-hIgG, 5) 180 kBq ^212^Pb-TCMC-chOI-1, or 6) 405 kBq ^212^Pb-TCMC-chOI-1 (10:1 TCMC:Ab ratio for both OI-1 and hIgG) with an antibody amount of 10 µg per mouse for groups 2, 4, and 6, and 5 µg for groups 3 and 5.

Tumor growth was visualized by bioluminescence imaging on day 43 using an IVIS Spectrum *in vivo* imaging system (PerkinElmer, Waltham, MA). For imaging, each mouse was injected IP with 0.2 ml of D-luciferin (Biosynth AG, Staad, Switzerland) dissolved in DPBS at a concentration of 20 mg/ml. The mice were anesthetized with sevoflurane and imaged 10 min after the luciferin injection.

Mice were observed throughout the study by monitoring their body weights (Fig. S1), activity level, posture, and IP tumor development every 2-3 days. All animals were euthanized by cervical dislocation at the study endpoint (day 52/53) when the tumor burden was high in the control group, in which the mice were close to the humane endpoint. The tumors were excised, and the total intraperitoneal tumor weight in each mouse was measured at the study endpoint.

Statistically significant differences between the resulting tumor weights were evaluated using the GraphPad software. One-way ANOVA was employed, and the obtained p-values were adjusted using Tukey’s method for multiple comparisons, with a significance threshold of P_adj_<0.05.

Tumor-free fraction (TFF) analysis was performed to evaluate treatment efficacy. Mice were considered tumor-free if their tumor weight was ≤0.020 g (lower limit of the scale was 0.020 g, controlled with a standard weight of 20.0 mg) at the end of the study period. The tumor-free fraction was calculated for each group by dividing the number of tumor-free mice by the total number of mice in each group. The statistical significance of the differences in TFF between the ^212^Pb-TCMC-chOI-1 treated groups and the control groups was determined using GraphPad software and two-sided Fisher’s Exact Test, with a significance threshold of p<0.05.

## Results

### Identification and validation of the target of OI-1 and affinity measurements

The mOI-1 antibody was screened for binding against a microarray of transfected HEK293 cells expressing 6100 individual full-length human plasma membrane proteins, cell-surface-tethered human secreted proteins, and 396 human heterodimers. The pre-screen confirmed minimal non-specific background binding of the mOI-1 antibody to untransfected HEK293 cells and no binding to any of the spotted targets when added before fixation, ensuring that interactions with target transfected cells in the library and confirmation screens would be distinguishable from untransfected cells. The following library screen identified 36 library interactions with spot intensities ranging from weak to strong. In the confirmation screen performed on live cells before fixation, 32 of the 36 library interactions were reproducibly observed using the mOI-1 antibody. Twenty-five of the 36 library interactions were classified as non-specific because they were bound by the mOI-1 antibody and by at least one of the negative control antibodies. These non-specific interactions included Fc gamma receptors.

After excluding non-reproducible and non-specific interactions, seven interactions remained. The mOI-1 antibody specifically interacted with PTK7 (isoform 4, medium intensity; isoform 1, strong intensity), TEK (very weak and weak intensity), ASGR2 (isoform 3, strong intensity; isoform 1, medium intensity), GH1 (weak/medium and medium intensity), and THPO (weak/medium intensity) (Table 1). Although classified as non-specific, the mOI-1 antibody showed an interaction with ROBO2 that was of greater intensity with the mOI-1 than with the controls (Table 1).

**Table 1.** Specific protein interactions of the mOI-1 antibody identified by screening a library of cell surface expressed proteins. Specific tein interactions were observed using the mOI-1 antibody in fixed and live cell confirmation screens. Proteins are listed according to their pective isoforms and their interaction intensities. Green and blue color indicate specific interactions, medium/strong or weak respectively. rple color indicates non-specific interaction though with greater intensity with mOI-1 than control antibodies. For flow cytometry fold change median fluorescence intensity (fc MFI) compared to ZsGreen only transfected HEK293 is shown (N=2). p: Repetition NI: No interaction. NT: Not tested.

| Gene Id | Protein Name | Library screen<br>(live cells, before<br>fixation) 36 interactions<br>in total |  | 1<sup>st</sup> Confirmation screen<br>(live cells, before<br>fixation) 7+1<br>interactions in total |  | 2<sup>nd</sup><br>Confirmation<br>screen<br>(absence of<br>fixation) | Verification screen<br>(flow cytometry) fc<br>MFI |
| --- | --- | --- | --- | --- | --- | --- | --- |
|  |  | Rep 1 | Rep 2 | Rep 1 | Rep 2 |  |  |
| PTK7 | Inactive tyrosine-protein<br>kinase 7 (Isoform 1) | strong | strong | strong | strong | strong | 10.0-11.7 (strong) |
|  | Inactive tyrosine-protein<br>kinase 7 (Isoform 4) | medium | medium | medium | weak/med | weak/med | 3.5 - 3.9 (weak) |
| TEK | Angiopoietin-1 receptor | v.weak | NI | v.weak | weak | NI | NT |
| ASGR2 | Asialoglycoprotein receptor 2<br>(Isoform 1) | medium | medium | medium | medium | NI | 1.5 – 1.8 (NI) |
|  | Asialoglycoprotein receptor 2<br>(Isoform 3) | med/strong | med/strong | strong | strong | NI | 1.7 – 1.9 (NI) |
| GH1 | Somatotropin | weak | weak/med | weak/med | medium | NI | 1.2 (NI) |
| THPO | Thrombopoietin | medium | weak/med | weak/med | weak/med | NI | 1.2 (NI) |
| ROBO2 | Roundabout homolog 2 | strong | strong | med/strong | med/strong | medium | 3.5 – 4.1 (weak/med) |

In a second confirmation screen conducted on live cells without fixation, the mOI-1 antibody resulted in specific interactions with PTK7 isoforms 1 and 4 (which differ by only a few amino acids (34)), whereas interactions with TEK, ASGR2 (isoforms 1 and 3), GH1, and THPO were not observed (Table 1). This suggests that the interactions with these proteins may have been artifacts of the fixation process. The interaction with ROBO2, despite being non-specific, was again noted for its increased intensity over controls, warranting further investigation. Flow cytometry on live transfected HEK293 cells verified a strong interaction of mOI-1 with PTK7 isoform 1, weak interaction with PTK7 isoform 4, and weak to medium interaction with ROBO2 (Table 1).

SPR was employed to validate the target specificity of OI-1 and to perform affinity measurements for various antibody constructs, including mOI-1 and chOI-1. SPR analysis revealed that chOI-1 exhibited binding to PTK7 and no discernible interaction with ROBO2, whereas its binding to PTK7 was best fitted by a two-state analysis model (Table 2). This suggests that chOI-1 binding to PTK7 may involve a complex interaction mechanism as a typical 1-to-1 binding profile was not observed. Such a mechanism could involve conformational changes in PTK7 upon antibody binding, further highlighting the intricate nature of this molecular interaction. The K_D_ obtained for mOI-1 was identical to that obtained for chOI-1 when analyzed using the two-state model for PTK7.

**Table 2.** Affinity measurement of mAbs. K_D_ of TCMC-and non-TCMC-conjugated mAbs by surface plasmon resonance analysis of human PTK7 or ROBO2 proteins. K_D_: equilibrium dissociation constant. Two state reaction analysis was used for all interactions.

| Antibody | Antigen | $K_D$ (nM) |
| --- | --- | --- |
| chOI-1 | ROBO2 | No binding detected |
| | PTK7 | $63.1 \pm 31.1$ (n=3)* |
| mOI-1 |  | 47.8 (n=1)** |
| 5:1 TCMC:chOI-1 |  | 81.1 (n=1) |
| 10:1 TCMC:chOI-1 |  | 98.7 (n=1) |
| 20:1 TCMC:chOI-1 |  | 155 (n=1) |

\* Combination of single cycle kinetics and multicycle kinetics data
\*\* Chip coupled with anti-mouse IgG antibody

### Immunohistochemistry results with OI-1 on human primary ovarian cancer

Slide evaluation of the ovarian cancer tissue sections revealed positive membranous staining for both chOI-1 and mOI-1 antibodies in the epithelium and tumor cells (Table 3, Fig. 1). The staining pattern was consistent between the chimeric and murine versions of the antibody.

**Fig. 1.**
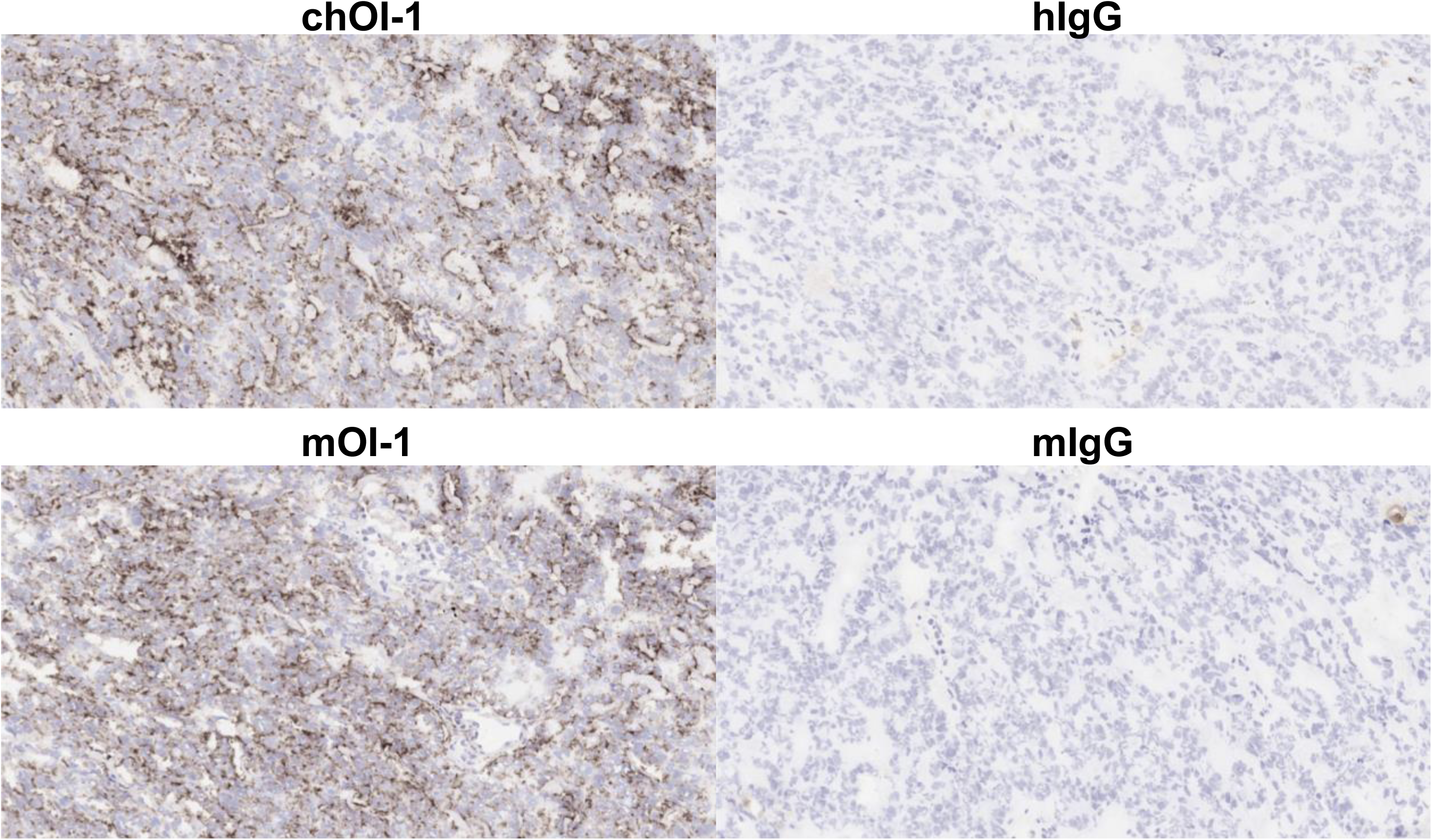
Immunohistochemical slides of primary ovarian cancer cells stained with OI-1. Slides were stained with 2.5 µg/mL mOI-1, chOI-1, mIgG or hIgG conjugated with Alexa 488.

**Table 3.** PTK7 expression in human primary ovarian cancer cells. Slide evaluation of ovarian cancer tissues using different concentrations (10, 2.5, and 0.625 μg/mL) of chOI-1 and mOI-1. Staining intensity was scored 0-4:0 (none), 1 (weak), 2 (mild), 3 (moderate), and 4 (high, potentially obscuring cellular structures). The staining frequency was graded as - (0%), + (≤25%), ++ (26-50%), +++ (51-75%), and ++++ (≥76%) for cells showing positive staining.

| Test Article | Concentration ($\mu\text{g/mL}$) | Staining Intensity | Staining Frequency |
| --- | --- | --- | --- |
| chOI-1 | 10 | 4 | +++ |
|  | 2.5 | 4 | +++ |
|  | 0.625 | 3 | +++ |
| mOI-1 | 10 | 4 | +++ |
|  | 2.5 | 3 | +++ |
|  | 0.625 | 3 | +++ |

### Cell surface expression and internalization of chOI-1 on SKOV-3-luc

Flow cytometry confirmed the cell surface expression of PTK7 in the SKOV-3-luc cancer cell line using various concentrations of chOI-1 antibody (Fig. 2A). A saturation binding experiment measured the equilibrium binding to PTK7 in SKOV-3-luc cells, determining the apparent K_D_^APP^ of 15.9 ± 2.8 nM based on rMFI data (Fig. 2B).

**Fig. 2.**
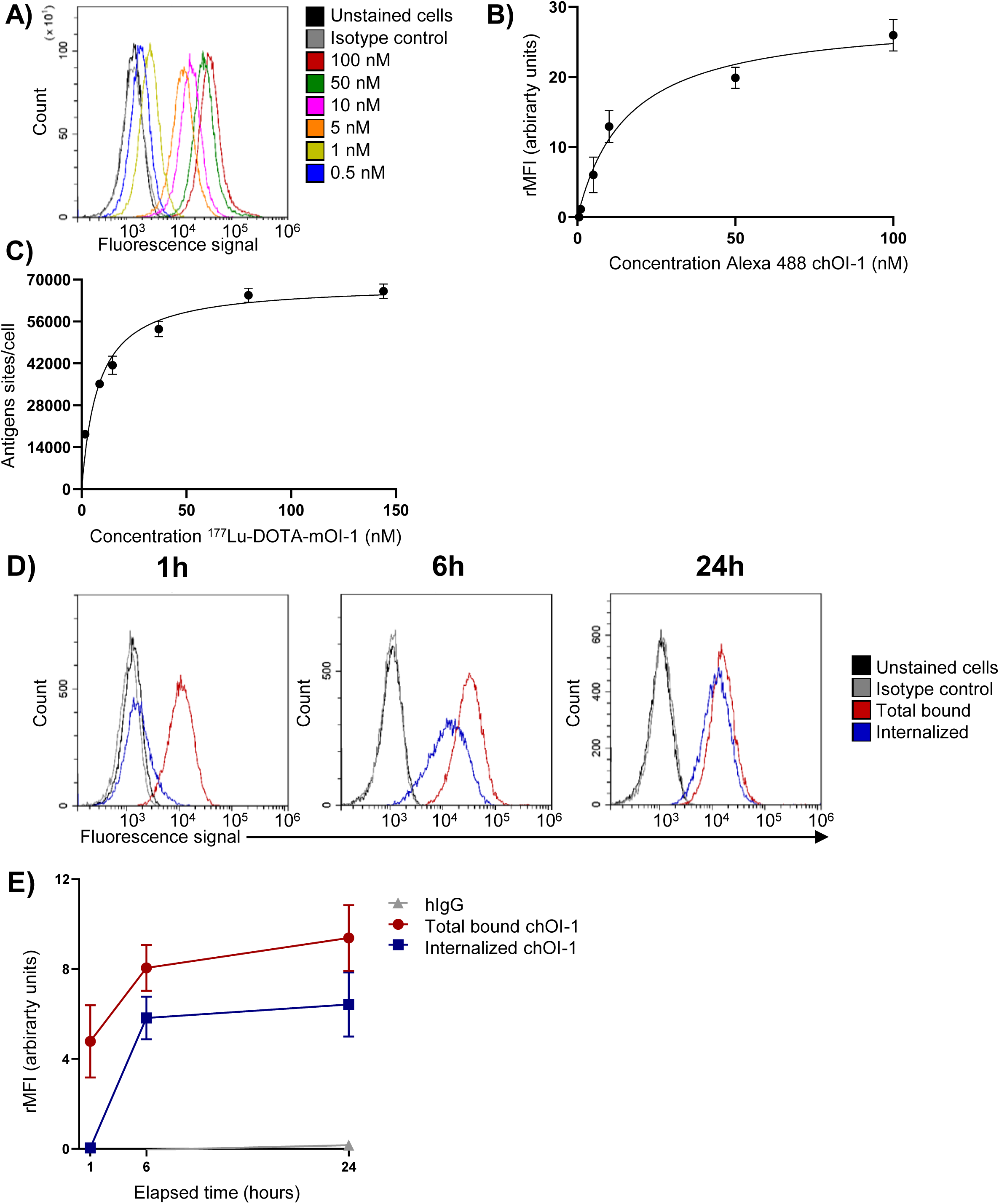
Binding and internalization of chOI-1 to PTK7 in SKOV-3-luc cells. Flow cytometry experiments were performed using Cytoflex, and only MitoTracker Red CMXRos and DRAQ5 double-positive cells were analyzed. A) Representative histograms showing the binding of chOI-1 Alexa 488 (0.5-100 nM) to SKOV-3-luc cells at 4°C for 30 min. Unstained cells and isotype controls served as negative controls B) Relative median fluorescence intensity (rMFI) analysis using a one-site binding model (Y = Bmax × X / (K_D_^APP^ + X)). For SKOV-3-luc cells, the apparent K_D_^APP^ for chOI-1 was 15.9 ± 2.8 nM. C) Saturation binding of ^177^Lu-DOTA-mOI-1 (1.6-144 nM) to SKOV-3-luc cells at 4°C for 65-95 minutes. Non-specific binding was determined by pre-incubation with excess unlabeled mAbs. The graph shows antigen sites/cell vs. ^177^Lu-DOTA-mOI-1 concentrations. A one-site binding model was used to determine K_D_^APP^. Points represent the mean ± SD of quadruplicate samples. D) Representative histograms showing internalization of chOI-1 Alexa 488 (0.25 µg) in SKOV-3-luc cells at 1, 6, and 24 h. The cells were incubated at 37°C, collected at specified time points, and either treated with trypsin (0.01 g/mL, 1, or 37°C) or left untreated. E) rMFI analysis of internalization over time. Points represent the mean of triplicate rMFI values ± SD from three independent experiments. The rMFI was calculated based on the MFI of unstained cells in each experiment.

Similarly, the saturation binding assay of ^177^Lu-DOTA-mOI-1 revealed a K_D_ of 7.9 ± 2.9 nM (n=2) (Fig. 2C). The maximum number of binding sites per cell (B_max_) was determined to be 68605 ± 4490. These results demonstrated that ^177^Lu-DOTA-mOI-1 binds specifically and with high affinity for SKOV-3-luc cells.

The internalization of the chOI-1 antibody in SKOV-3-luc cells was investigated using flow cytometry (Fig. 2E). After 1 h of incubation of SKOV-3-luc cells with chOI-1 Alexa 488, the surface-bound mAb was stripped using trypsin. The Alexa 488 signal was 2.6 ± 0.3% of the total signal detected in non-stripped cells (Table 4). At 6 h and 24 h, the signal increased to 71.8 ± 12.5% and 68.3 ± 9.9%, respectively (Table 4), indicating significant internalization of chOI-1 over a 6 hour time period followed by stabilization of internalization (Table 4). The rMFI data at the indicated time points showed a similar trend, with maximum internalization at 6 h (Fig. 2E). In contrast, the human isotype control Alexa 488 showed no internalization during this time period (Fig. 2E).

**Table 4.** Percentage internalization of Alexa 488 mAbs in Skov-3-luc cells. Skov-3-luc cells were incubated with 0.25 µg chOI-1 Alexa 488 for 1, 6, or 24 h. The cells were either left untreated (total signal) or exposed to trypsin to remove the surface-bound antibodies (internalized signal). Only live cells were included in this analysis. The relative fluorescence intensity (rMFI) of each sample was measured, and the percentage of internalized signal was calculated as the internalized rMFI signal (trypsin-stripped cells) compared with the total-bound rMFI (non-stripped cells). Data are representative of three independent experiments.

| Incubation time<br>(h) | Internalized<br>chOI-1<br>(%) |
| --- | --- |
| 1 | 2.6 ± 0.3 |
| 6 | 71.8 ± 12.5 |
| 24 | 68.3 ± 9.9 |

### Effect of molar ratios of TCMC during chelation reaction of mAb on antibody binding to PTK7

The effect of the molar ratio of TCMC during the chelation reaction of mAb on antibody binding to PTK7 was investigated. The IRF results for ^212^Pb-TCMC-chOI-1 demonstrated a clear inverse relationship between the TCMC-to-antibody ratio and IRF values (Table S1). As the TCMC-to-antibody ratio increased from 5:1 to 10:1 and 20:1, IRF decreased significantly (p<0.01). For the SKOV-3-luc cell line, the IRF values were 65.3 ± 4.9%, 53.5 ± 3.9%, and 43.3 ± 3.2% for the 5:1, 10:1, and 20:1 ratios, respectively (Table S1). Similarly, for the OHS cell line, IRF values were 59.6 ± 2.7%, 46.3 ± 8.2%, and 40.3 ± 1.5% at the same ratios (Table S1). This trend indicated that a higher molar ratio of TCMC negatively affected the ability of the antibody to bind to its target antigen.

Mass spectrometry was used to determine the number of TCMC attached to chOI-1 (Fig. S2). At a 5:1 ratio, most antibodies had 1-2 TCMC molecules conjugated to each mAb, with approximately 15% of chOI-1 not bound to TCMC. A 10:1 ratio resulted in 3-6 TCMC molecules attached per antibody, up to 7, with approximately 5% unbound chOI-1. At a 20:1 ratio, most antibodies contained up to seven TCMC molecules, with some reaching concentrations as high as 10.

SPR analysis was used to determine the influence of an increased number of conjugated TCMC in chOI-1 on K_D_ measurements using human PTK7. The K_D_ values for different TCMC-conjugated antibodies showed a decreasing trend in affinity with increasing chelator-to-antibody ratios (Table 2). Notably, the 5:1 and 10:1 TCMC:Ab ratio conjugates maintained affinities within 2-fold of the unconjugated chOI-1 (Table 2), whereas the 20:1 TCMC conjugate showed approximately 2.5-fold lower affinity. An overall decrease in signal was observed with increasing TCMC conjugation, suggesting a reduction in the proportion of antibody molecules capable of binding to the antigen. Based on these data, only 5-and 10-fold TCMC ratios were used in the subsequent studies.

### Biodistribution of ^177^Lu-DOTA-mOI-1 in SKOV-3-luc Xenograft Models

*In vivo* biodistribution studies were conducted in mice bearing subcutaneous SKOV-3-luc xenografts to evaluate the tumor-targeting capabilities of ^177^Lu-DOTA-mOI-1 following intravenous administration. The percentage of injected dose per gram of tissue (% ID/g) was measured over a 7-day period in the tumors and normal organs (Fig. 3, Table S2). Tumor uptake progressed significantly from A at day 1 to B at day 7 (p_adj_<0.03). In contrast, most normal tissues show decreased or stable uptake over time. The blood retention of ^177^Lu-DOTA-mOI-1 decreased significantly from 15.4 ± 4.4% ID/g on day 1 to 4.5 ± 1.3% ID/g by day 7 (p_adj_ <0.003). In addition, the muscle and small bowel showed a significant decrease in uptake from days 1 to 7 (p_adj_<0.03). A significant difference was observed between blood and tumor uptake on days 3, 4, and 7 (p_adj_<0.01). All other normal tissues showed significantly less uptake than the tumor (p_adj_<0.004).

**Fig. 3.**
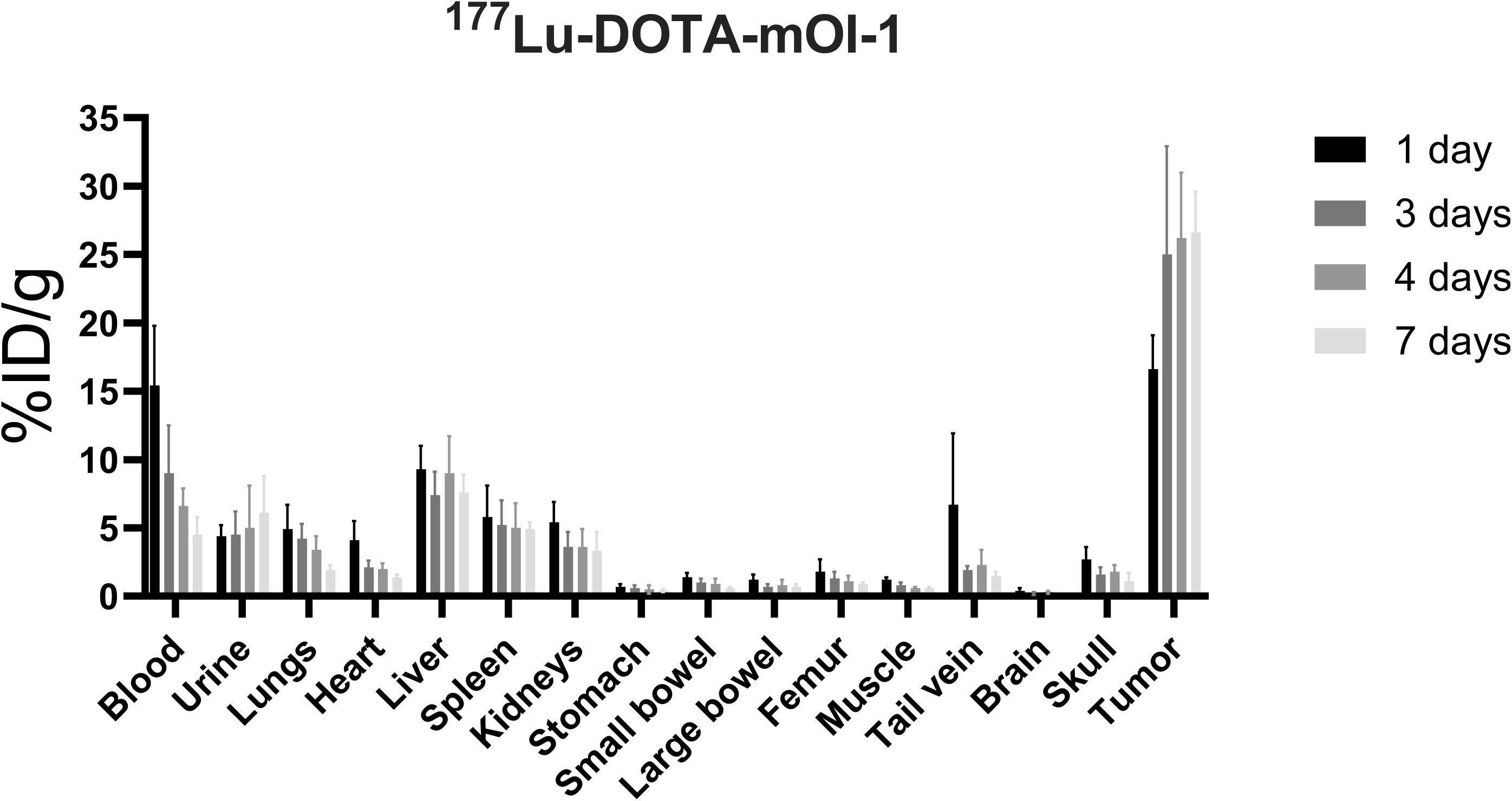
Biodistribution of ^177^Lu-DOTA-mOI-1 in mice bearing SKOV-3-luc xenografts. The percentage of injected dose per gram (% ID/g) was measured in nude mice subcutaneously inoculated with SKOV-3-luc tumor cells. Mice were intravenously injected with 450 kBq ^177^Lu-DOTA-mOI-1 35 days after cell inoculation. Tissue samples were collected at various time points: 1, 3, 4, and 7-days post injection. Bars represent the mean % ID/g and error bars indicate the standard deviation; the study included four mice per time point for ^177^Lu-DOTA-mOI-1.

### Therapeutic efficacy of ^212^Pb-TCMC-chOI-1 in intraperitoneal ovarian cancer

The administration of ^212^Pb-TCMC-chOI-1 demonstrated significant therapeutic efficacy in mice with intraperitoneal ovarian cancer xenografts, with a marked reduction in tumor burden observed in the ^212^Pb-TCMC-chOI-1 groups compared to the non-radioactive groups at the study endpoint (day 52/53). TFF analysis showed that the group treated with 180 kBq of ^212^Pb-TCMC-chOI-1 achieved a TFF of 100%, indicating complete tumor eradication, while the 405 kBq group achieved a TFF of 87.5% (Table S3). In contrast, the ^212^Pb-TCMC-hIgG-treated groups had a TFF of only 25%. Statistical analysis confirmed these results, revealing significantly higher TFF in the ^212^Pb-TCMC-chOI-1-treated groups than in the control groups, with the 180 kBq group showing a significant difference in TFF compared to both ^212^Pb-TCMC-hIgG groups (p=0.007), which was also the case for the 405 kBq group (p=0.04). Furthermore, tumor weight measurements demonstrated significant reductions in the ^212^Pb-TCMC-chOI-1 groups compared to the non-radioactive groups, reinforcing the anti-tumor effects of ^212^Pb-TCMC-chOI-1 (Fig. 4). While both ^212^Pb-TCMC-hIgG groups also showed a significant reduction in tumor weight, the effect was less pronounced.

**Fig. 4.**
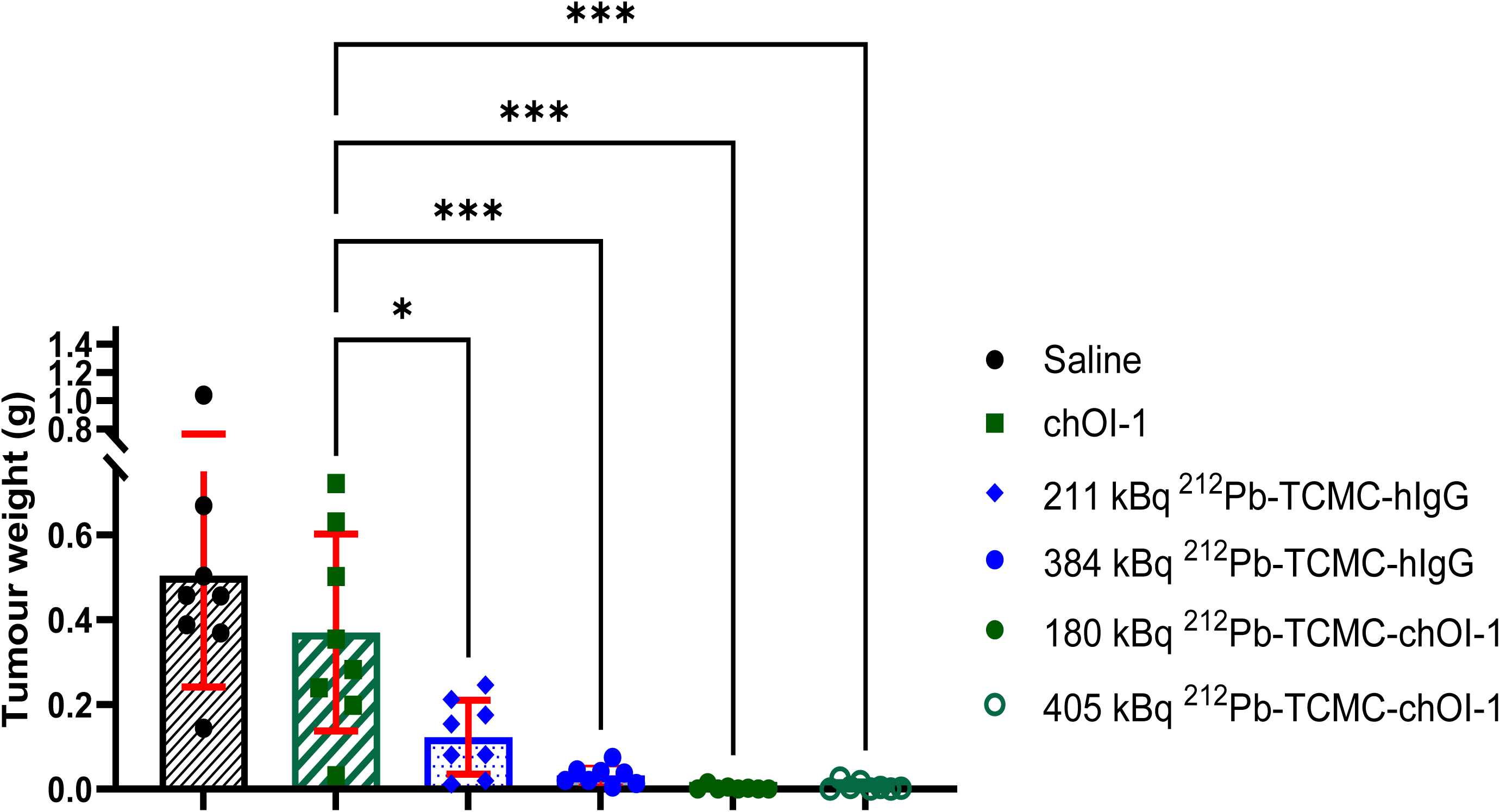
Anti-tumor effect of ^212^Pb-TCMC-chOI-1. The intraperitoneal (IP) mean ± standard deviation tumor weight after euthanizing nude mice on day 52/53 injected IP with SKOV-3-luc cells on day zero and treated IP on day 3 with 1) saline, 2) chOI-1, 3) 211 kBq ^212^Pb-TCMC-hIgG; 4) 384 kBq ^212^Pb-TCMC-hIgG, 5) 180 kBq ^212^Pb-TCMC-chOI-1, or 6) 405 kBq ^212^Pb-TCMC-chOI-1, with an antibody concentration of 10 µg per mouse for groups 2, 4, and 6 and 5 µg for groups 3 and 5.Eight mice were used for each group. P-values were calculated using a one-way ANOVA test with correction for multiple comparisons using Tukey’s method; *p_adj_<0.02 and ***p_adj_<0.002. All radioactive treatment groups (3–6) showed similarly significant reductions in tumor weight compared to the saline group, as the chOI-1 group.

Further supporting these results, bioluminescence imaging on day 43 revealed a markedly lower tumor burden in the ^212^Pb-TCMC-chOI-1 groups compared to the ^212^Pb-TCMC-hIgG groups (Fig. 5). These comprehensive analyses underscore the therapeutic efficacy of ^212^Pb-TCMC-chOI-1 in reducing the tumor burden and achieving tumor-free outcomes.

**Fig. 5.**
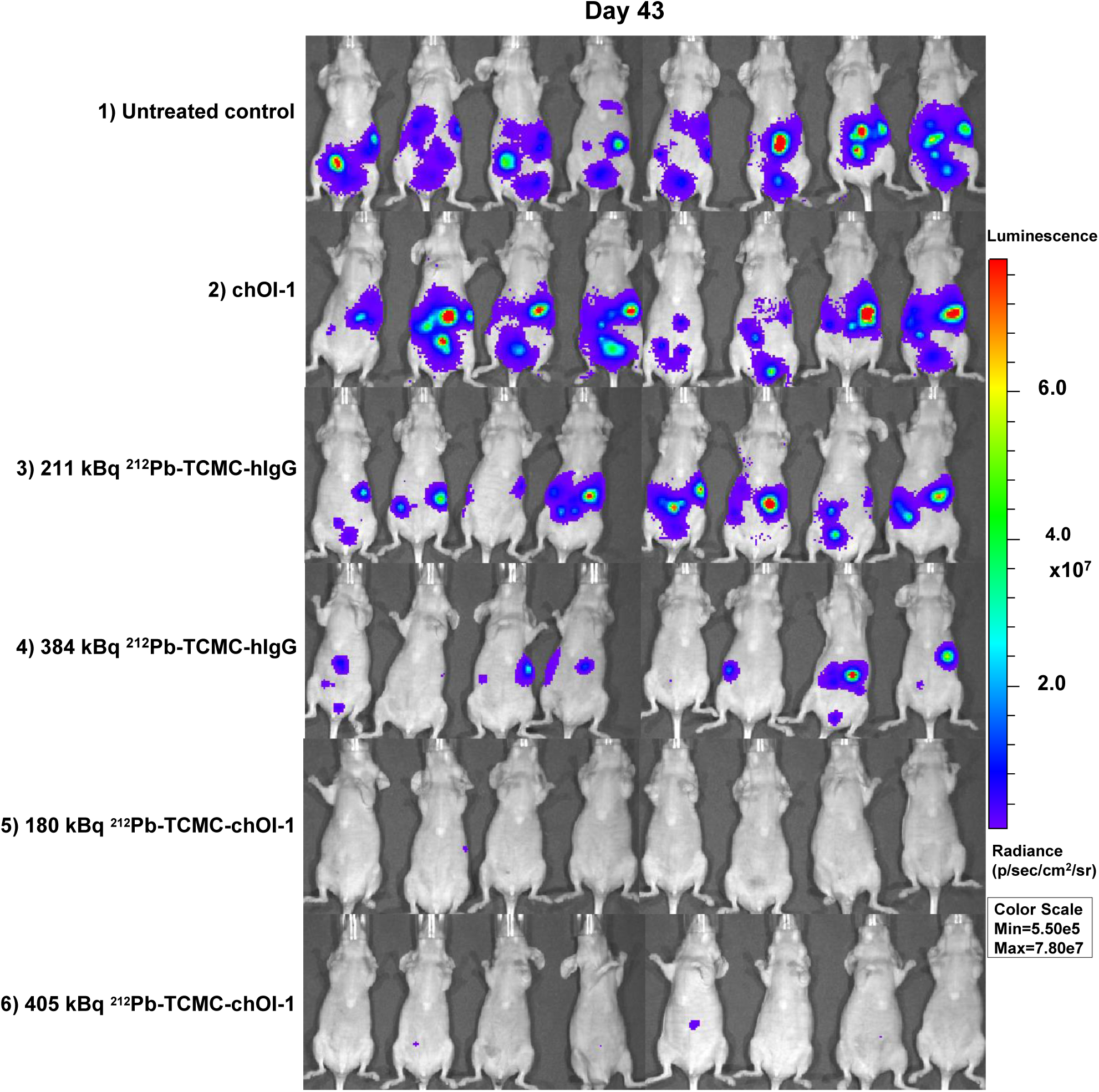
Tumor visualization in treated mice. Tumor growth was visualized by bioluminescence imaging in the different therapy groups on day 43 after SKOV-3-luc inoculation. For imaging, each mouse was injected intraperitoneally with d-luciferin. The mice were anesthetized with sevoflurane and imaged 10 min after the luciferin injection. Eight mice were used for each group.

## Discussion

The current study is to our knowledge the first to demonstrate the potential of targeting the PTK7 receptor for tumor selective delivery and effect of radionuclide therapy. The human PTK7 antigen was identified and validated as the target of the murine OI-1 antibody, and conversion to chimeric OI-1 had no impact on binding. Our results showed weaker binding to PTK7 isoform 4. While the reasons for this are unclear, isoform 4 is less expressed, not well characterized, and structurally similar to isoform 1, although it is slightly shorter (1014 vs. 1070 amino acids) (34). The strong binding of OI-1 to isoform 1, as confirmed by SPR assays, supports its relevance in further investigations. Additionally, the lack of ROBO2 binding in our validation screen confirmed the specificity of OI-1 for PTK7. The OI-1 antibody variants showed binding affinities of 47.8-63.1 nM to human PTK7 in SPR analysis, which is lower than those reported for other PTK7-targeting antibodies in similar assays, cofetuzumab (0.7-1.4 nM) (1) and mAb13 (0.06 nM) (6). While lower affinity might reduce the immediate binding strength, it can enhance tissue penetration and distribution within tumors (35).

Immunohistochemical analysis confirmed that PTK7 effectively identified human primary ovarian cancer tissues, consistent with the findings of Kong et al., Damelin et al., and Xu et al., who also reported minimal expression in normal ovarian tissues (1, 6, 8). The role of PTK7 in ovarian cancer has been highlighted as complex by Wang et al., who reported lower PTK7 expression in epithelial ovarian carcinomas than in benign tumors, indicating a nuanced relationship (4). However, in a clinical phase 1 trial investigating a cofetuzumab–drug conjugate in ovarian cancer (NCT02222922), patients were not preselected based on PTK7 expression levels (7). In contrast, patients with triple-negative breast cancer and non-small cell lung cancer were screened and required to have moderate-to-high PTK7 expression. Post-treatment assessments determined that more than 80 percent of patients with ovarian cancer were positive for PTK7 expression (7). Collectively, these findings underscore the potential of PTK7 as a target for ovarian cancer therapy.

Our *in vitro* studies using the human ovarian cancer cell line SKOV-3-luc confirmed PTK7 expression and internalization with properties similar to those reported by Kong et al. for their anti-PTK7 mAb (6).

In drug-loaded therapies including ARCs, optimizing the chelator-to-antibody ratio (CAR) is important for balancing the radiolabeling efficiency and maintaining the functional integrity of the antibody. In our study, we found that a molar ratio of 5:1 to 10:1 TCMC:Ab during the chelation reaction provided suitable antibody-binding properties for ^212^Pb-TCMC-mAb conjugates. Our findings align with those of previous research on DOTA-based conjugation (a chelator structurally similar to TCMC (36, 37)). Although higher CARs can increase the specific activity by attaching more radionuclides per antibody, this may come at the cost of reduced binding affinity due to steric hindrance or conformational changes. For ^212^Pb-based therapies, the specific activities reported in the literature typically range from 37 to 370 MBq/mg (38), depending on the experimental setup. In our therapy study, we used a specific activity of approximately 50 MBq/mg, which is within this range.

Our study evaluated the systemic biodistribution of mOI-1 in xenograft mice bearing subcutaneous tumors, using ^177^Lu (t_1/2_= 6.7 d) to align with the pharmacokinetics of the antibody and accurately assess tumor targeting over 1 to 7 days. The longer half-life of ^177^Lu facilitated an evaluation of the anti-PTK7 antibody’s tumor-targeting capabilities. The percentage of injected dose per gram was consistently higher in tumors than in any normal organ. This biodistribution profile demonstrated effective and specific tumor targeting, supporting potential applications in radioimmunotherapy using the SKOV-3-luc model and confirming its accumulation in subcutaneous ovarian cancer tumors via systemic IV administration.

Regarding normal tissue binding, PTK7 is expressed at low levels in some normal tissues (1), which should be considered when translating these findings from mice to humans. Additionally, the high protein sequence similarity between human and murine PTK7, 93% identity, and 97% positivity (NP_002812.2 vs. ID Q8BKG3.1), supports the likelihood of species cross-reactivity of the OI-1 antibody to murine PTK7, further validating the use of this model system in preclinical studies.

While IV administration was used to assess selective tumor targeting and the systemic distribution of ^177^Lu-DOTA-mOI-1, we propose that IP administration could be highly relevant for ^212^Pb-mAb therapy in ovarian cancer with peritoneal dissemination. IP injections provide direct access to peritoneal metastases, achieving higher local concentrations of ARC at the tumor site than IV delivery (39). This method aligns the physical half-life of ^212^Pb with the biological half-life of mAbs, thereby enhancing the therapeutic concentration at the tumor site and potentially minimizing the radiation exposure to normal organs (40).

The therapeutic potential of ^212^Pb-TCMC-chOI-1 was clearly demonstrated in a mouse model with intraperitoneal ovarian cancer xenografts, particularly at lower doses, as indicated by the high TFF. The increased efficacy of ^212^Pb-TCMC-chOI-1 compared with non-specific ^212^Pb-TCMC-hIgG further underscores the importance of targeted therapy for achieving better tumor control.

While our study demonstrated significant therapeutic potential for ^212^Pb-TCMC-chOI-1 in treating intraperitoneal ovarian cancer, there are some limitations to consider. The comparison of tumor masses at a single time point may have limited our ability to observe the full extent of the treatment’s effects. A longer observation period could have revealed a more pronounced difference between the ^212^Pb-TCMC-hIgG and ^212^Pb-TCMC-chOI-1 groups, as allowing the mice to live longer might have provided clearer evidence of therapeutic efficacy. Although our biodistribution analysis included normal tissue evaluation, we did not conduct toxicity assessments, and further investigations are needed to fully assess its safety. Future preclinical studies should focus on evaluating the therapeutic effects over longer time periods (e.g., through survival studies), conducting toxicity assessments, and optimizing treatment parameters.

Our findings show the potential of the PTK7 targeting OI-1 antibody as an ARC, with this first proof-of-principle study of its therapeutic effect in combination with a short-lived radionuclide for intraperitoneal treatment of ovarian cancer. Targeted antibody delivery combined with high-LET alpha emissions from ^212^Pb’s daughter ^212^Bi shows promise for localized treatment, exemplified by intraperitoneal administration. Given the PTK7 expression in various malignancies, including triple-negative breast cancer, non-small-cell lung cancer (41), and bladder cancer (42), these preliminary results suggest the possibility of expanding this therapeutic approach to other PTK7-expressing cancers.

## Supplementary Information

The online version contains supplementary material available.

**Additional file 1: Fig. S1.** Body weight monitoring of animals in the ^212^Pb-chOI-1 study efficacy study. **Fig. S2.** Mass spectrometry analysis of TCMC-conjugated chOI-1 with different TCMC:mAb ratios during the chelating reaction. **Table S1.** Immunoreactive affinity of antibody-radionuclide conjugate (ARC). **Table S2.** Biodistribution (%ID/g) of 177Lu-DOTA-mOI-1 in mice bearing SKOV-3-luc xenografts. **Table S3.** Tumor weight of animals in the ^212^Pb-chOI-1 study efficacy study.

## Declarations

### Ethics approval and consent to participate

All experimental procedures involving animals were approved by the Institutional Committee on Research Animal Care (Department of Comparative Medicine, Oslo University Hospital, Norway) and the Norwegian Food Safety Authority. The animal experiments were approved under project number 23766 on 21 September 2020. The animals were treated in accordance with institutional, national, and EU regulations on the use of research animals.

### Consent for publication

Not applicable.

### Availability of data and materials

The data that support the findings of this study are available from Oncoinvent ASA, but restrictions apply to the availability of these data, which were used under license for the current study, and so are not publicly available. Data are however available from the corresponding author at upon reasonable request and with permission of Oncoinvent ASA.

### Authors’ contributions

Conceptualization, K.L., S.W., R.G.L., S.M.D., M.M.M. and T.B.B.; methodology, K.L., S.W., S.M.D., R.G.L., and M.M.M.; validation, K.L., S.W., S.M.D., M.M.M. R.G.L. and.; formal analysis, K.L., S.W., B.H., and S.M.D; investigation, K.L., S.W., S.M.D., M.M.M., B.H., R.G.L. and S.W.; resources, T.B.B.; data curation, T.B.B.; writing—original draft preparation, K.L..; writing—review and editing, K.L., S.M.D., R.G.L., A.J., T.B.B., and S.W.; visualization, K.L.; supervision, T.B.B., A.J., M.M.M. and S.M.D.; project administration, T.B.B.; funding acquisition, T.B.B.. All authors have read and agreed to the published version of the manuscript.

#### Acknowledgements

We would like to thank the staff at Oncoinvent ASA, including Zeljka Raskovic-Lovre for assistance with setting up the ^212^Pb generators, and Carina Hinrichs and Anna Amalie Zickfeldt Lade for their help with cell culturing.

