## Supplementary information for "Preclinical Evaluation of PTK7-Targeted Radionuclide Therapy"

**Table S1. Immunoreactive affinity of antibody-radionuclide conjugate (ARC).** The average radiochemical purity (RCP) was >98% for all RIC, and the average immunoreactive fraction (IRF) was expressed as the cell bound percentage of RIC ± SD.
SD, standard deviation
n, number of samples
NM, not measured

| ARC | Molar ratio of chelator to mAb in chelation reaction | Average RCP of RIC (%) ± SD | Average IRF (%) ± SD of RIC | |
|---|---|---|---|---|
| | | | SKOV-3-luc | OHS |
| $^{177}$Lu-DOTA-mOI-1 | 5:1 | 99.4 ± 0.5 (n=8) | NM | 36.7 ± 4.5 (n=3) |
| $^{212}$Pb-TCMC-mOI-1 | 5:1 | 99.0 ± 0.9 (n=6) | 44.3 ± 2.1 (n=3) | 42.3 ± 1.7 (n=4) |
| $^{212}$Pb-TCMC-chOI-1 | 5:1 | 98.3 ± 1.0 (n=11) | 65.3 ± 4.9 (n=4) | 59.6 ± 2.7 (n=5) |
| | 10:1 | 98.3 ± 1.6 (n=16) | 53.5 ± 3.9 (n=4) | 46.3 ± 8.2 (n=11) |
| | 20:1 | 99.3 ± 1.0 (n=4) | 43.3 ± 3.2 (n=3) | 40.3 ± 1.5 (n=3) |

**Table S2. Biodistribution (%ID/g) of $^{177}$Lu-DOTA-mOI-1 in mice bearing SKOV-3-luc xenografts.** The percentage of the injected dose per gram (% ID/g) was assessed in nude mice that had been subcutaneously inoculated with SKOV-3-luc tumor cells. The mice received intravenous injections of 450 kBq of $^{177}$Lu-DOTA-mOI-1, administered 35 days after tumor cell inoculation. Tissue samples were collected at different intervals: for $^{177}$Lu-DOTA-mOI-1, samples were taken at 1, 3, 4, and 7 days after injection. % ID/g is presented together with standard deviation. n=number of animals.

| | **$^{177}$Lu-DOTA-mOI-1** | | | |
|---|---|---|---|---|
| **Time point (days)** | **1 (n=4)** | **3 (n=4)** | **4 (n=4)** | **7 (n=4)** |
| Blood | 15.4 ± 4.4 | 9.0 ± 3.5 | 6.6 ± 1.3 | 4.5 ± 1.3 |
| Urine | 4.4 ± 0.8 | 4.5 ± 1.7 | 5.0 ± 3.1 | 6.1 ± 2.7 |
| Lung | 4.9 ± 1.8 | 4.2 ± 1.1 | 3.4 ± 1.0 | 1.9 ± 0.4 |
| Heart | 4.1 ± 1.4 | 2.1 ± 0.5 | 2.0 ± 0.4 | 1.4 ± 0.2 |
| Liver | 9.3 ± 1.7 | 7.4 ± 1.7 | 9.0 ± 2.7 | 7.6 ± 1.3 |
| Spleen | 5.8 ± 2.3 | 5.2 ± 1.8 | 5.0 ± 1.8 | 4.9 ± 0.5 |
| Kidney | 5.4 ± 1.5 | 3.6 ± 1.1 | 3.6 ± 1.3 | 3.3 ± 1.4 |
| Stomach | 0.7 ± 0.2 | 0.6 ± 0.2 | 0.5 ± 0.3 | 0.4 ± 0.1 |
| Small bowel | 1.4 ± 0.3 | 1.0 ± 0.3 | 0.9 ± 0.4 | 0.6 ± 0.1 |
| Large bowel | 1.2 ± 0.4 | 0.7 ± 0.2 | 0.8 ± 0.4 | 0.7 ± 0.2 |
| Femur | 1.8 ± 0.9 | 1.3 ± .06 | 1.1 ± 0.4 | 0.9 ± 0.1 |
| Muscle | 1.2 ± 0.2 | 0.8 ± 0.3 | 0.6 ± 0.1 | 0.6 ± 0.1 |
| Tail vein | 6.7 ± 5.2 | 1.9 ± 0.3 | 2.3 ± 1.1 | 1.5 ± 0.3 |
| Brain | 0.4 ± 0.2 | 0.2 ± 0.1 | 0.3 ± 0.1 | 0.1 ± 0.0 |
| Scull | 2.7 ± 0.9 | 1.6 ± .05 | 1.8 ± 0.5 | 1.1 ± 0.6 |
| Tumor | 16.6 ± 2.5 | 25.0 ± 7.9 | 26.2 ± 4.8 | 26.6 ± 3.0 |

**Table S3. Tumor weight of animals in the $^{212}$Pb-TCMC-chOI-1 study efficacy study.** Tumor weight in gram of n=8 nude mice per group is presented as individual weight for every mouse. Nude mice were inoculated intraperitoneally with SKOV-3-luc cells on day zero and treated intraperitoneally on day 3 with : 1) saline, 2) chOI-1, 3) 211 kBq $^{212}$Pb-TCMC-hIgG, 4) 384 kBq $^{212}$Pb-TCMC-hIgG, 5) 180 kBq $^{212}$Pb-TCMC-chOI-1, or 6) 405 kBq $^{212}$Pb-TCMC-chOI-1, with an antibody concentration of 10 µg per mouse for groups 2, 4, and 6, and 5 µg for groups 3 and 5. Mice were considered tumor-free if their tumor weight ≤0.020 g. TFF=Tumor free fraction

| Saline | chOI-1 | 211 kBq $^{212}$Pb-TCMC-hIgG | 384 kBq $^{212}$Pb-TCMC-hIgG | 180 kBq $^{212}$Pb-TCMC-chOI-1 | 405 kBq $^{212}$Pb-TCMC-chOI-1 |
|---|---|---|---|---|---|
| 0.503 | 0.282 | 0.154 | 0.021 | 0.005* | 0.018* |
| 0.369 | 0.354 | 0.175 | 0.013* | 0* | 0.004* |
| 0.669 | 0.630 | 0.012* | 0.038 | 0* | 0.026 |
| 0.144 | 0.502 | 0.080 | 0.005* | 0.002* | 0.002* |
| 0.388 | 0.721 | 0.212 | 0.045 | 0.015* | 0.004* |
| 1.040 | 0.239 | 0.081 | 0.075 | 0* | 0* |
| 0.456 | 0.198 | 0.246 | 0.042 | 0* | 0* |
| 0.457 | 0.032 | 0.020* | 0.021 | 0* | 0* |
| TFF: 0 % | TFF: 0 % | TFF: 25 % | TFF: 25 % | TFF: 100 % | TFF: 87.5 % |

*Considered tumor free

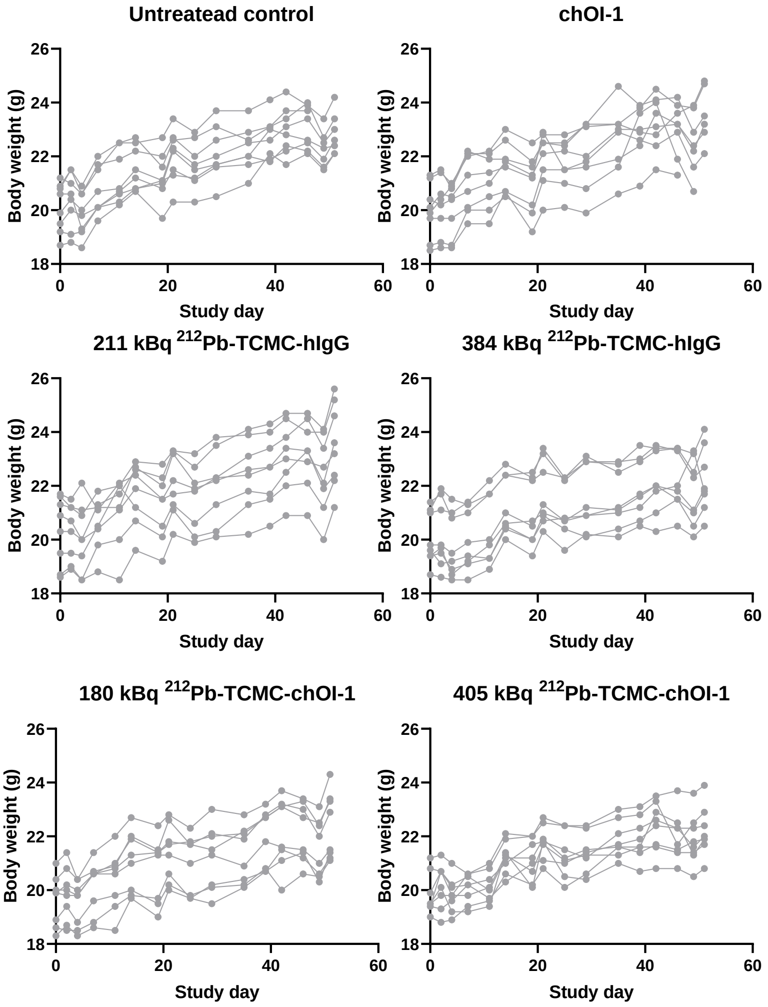


**Fig S1. Body weight monitoring of animals in the $^{212}$Pb-TCMC-chOI-1 study efficacy study.** Body weight of n=8 nude mice per group is presented as individual weight for every mouse. Nude mice were inoculated intraperitoneally with SKOV-3-luc cells on day zero and treated intraperitoneally on day 3 with : 1) saline, 2) chOI-1, 3) 211 kBq $^{212}$Pb-TCMC-hIgG, 4) 384 kBq $^{212}$Pb-TCMC-hIgG, 5) 180 kBq $^{212}$Pb-TCMC-chOI-1, or 6) 405 kBq $^{212}$Pb-TCMC-chOI-1, with an antibody concentration of 10 µg per mouse for groups 2, 4, and 6, and 5 µg for groups 3 and 5.

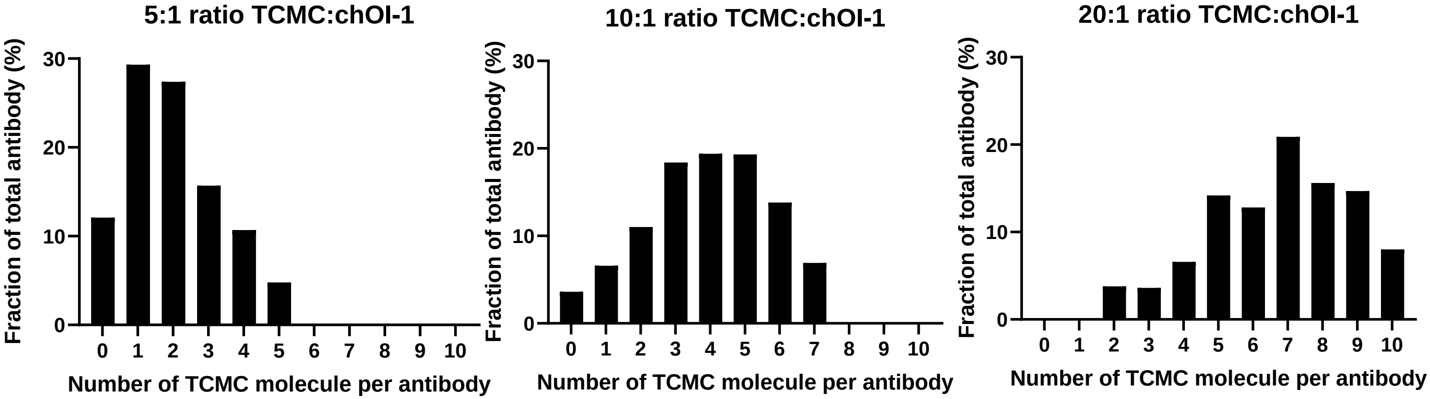


**Fig. S2. Mass spectrometry analysis of TCMC-conjugated chOI-1 with different TCMC:mAb ratios during the chelating reaction**. The number of TCMC molecules chelated to chOI-1 increased as the molar ratio of TCMC:mAb during the chelating reaction increased, as shown here for the molar ratios of 5:1, 10:1, and 20:1.
